# Protein identification with a nanopore and a binary alphabet

**DOI:** 10.1101/119313

**Authors:** G. Sampath

## Abstract

Protein sequences are recoded with a binary alphabet obtained by dividing the 20 amino acids into two subsets based on volume. A protein is identified from subsequences by database search. Computations on the *Helicobacter pylori* proteome show that over 93% of binary subsequences of length 20 are correct at a confidence level exceeding 90%. Over 98% of the proteins can be identified, most have multiple identifiers so the false detection rate is low. Binary sequences of unbroken protein molecules can be obtained with a nanopore from current blockade levels proportional to residue volume; only two levels, rather than 20, need be measured to determine a residue’s subset. This procedure can be translated into practice with a sub-nanopore that can measure residue volumes with ~0.07 nm^3^ resolution as shown in a recent publication. The high detector bandwidth required by the high speed of a translocating molecule can be reduced more than tenfold with an averaging technique, the resulting decrease in the identification rate is only 10%. Averaging also mitigates the homopolymer problem due to identical successive blockade levels. The proposed method is a proteolysis-free single-molecule method that can identify arbitrary proteins in a proteome rather than specific ones. This approach to protein identification also works if residue mass is used instead of mass; again over 98% of the proteins are identified by binary subsequences of length 20. The possibility of using this in mass spectrometry studies of proteins, in particular those with post-translational modifications, is under investigation.

## 1. Introduction

Sequencing/identification of peptides/proteins is currently based on Edman degradation, gel electrophoresis, or mass spectrometry (MS) [1–3]; it is usually done in the bulk and often followed by database search [3]. In comparison with DNA sequencing, for which several NGS (next generation sequencing) technologies are currently available [4], protein sequencing is more difficult, as it has to discriminate among 20 amino acids, as opposed to four bases with DNA. It also has to work with the available sample, as there is no amplification technique for proteins comparable to PCR for DNA [2].

Compared to protein sequencing, protein identification is simpler because it only requires finding a partial sequence and then searching through a protein sequence database to find the protein that is uniquely identified by it. Recently single-molecule methods based on proteolysis and optical or other labeling of selected residues have been proposed. In one of them a protein is cleaved into peptide fragments, pinned to a substrate, and selectively labeled [5]. The labeled residues are detected optically and a partial sequence obtained.

In contrast with these methods nanopores provide a single-molecule electrical alternative that does not require proteolysis, analyte immobilization, or labeling of any kind [6]. While their use in sequencing DNA is becoming commonplace [7], nanopore-based protein sequencing may seem like a distant prospect. Recent reviews of nanopore-based protein studies are available [8–11]. Examples of work in the area include experimental studies involving recognition tunneling [12] and use of a sub-nanometer diameter nanopore [13]. A major limiting factor has been the pore current resolution needed to discriminate among 20 different amino acids; at present this appears to be out of reach. There has been some success in nanopore-based studies of other aspects of proteins such as folding/conformation [14,15] and recognition of specific proteins or their variants [16–19].

### 1.1 The present work

Here it is shown by computation that proteins in a proteome can be uniquely identified from subsequences of primary sequences by using a binary code derived from a division of the amino acids into two sets based on published volume data [20]. Comprehensive computations on a sample proteome (*Helicobacter pylori*) show that the codes of subsequences 20 residues long are correct at a better than 90% confidence level, and that over 98% of the proteins in the proteome can be identified by searching for the subsequences in the binary-coded proteome database. With a nanopore this means that a pore current resolution that can discriminate between the two subsets mentioned above is sufficient. (Incidentally a binary alphabet has been used recently in DNA sequencing with a solid-state nanopore. In this approach a nucleotide is encoded with a predesigned oligonucleotide and two fluorescent label types to represent the four bases [21].)

The scheme described here can be translated into practice with existing technology (nanopore, electronic, database); an easy-to-use hand-held device similar to the MinION in genome sequencing [22] can be designed and implemented. Unlike most nanopore-based protein identification methods, the one presented here can identify arbitrary proteins in a proteome rather than specific individual targets [16–19]. As such it can also be extended to the analysis of protein mixtures (see Section 3.4).

## 2. Protein identification with a nanopore

An electrolytic cell has a membrane dividing two chambers (*cis* and *trans*) containing an electrolyte. A potential difference applied across the membrane leads to an ionic current through the pore from *cis* to *trans*. An analyte molecule (such as a polymer like DNA or protein) introduced into *cis* translocates through the pore to *trans* and causes a current blockade. A monomer may be identified from the blockade level, which may be specific to one or more contiguous monomers based on some physical or chemical property such as volume, charge, diffusion constant, etc. With proteins no enzymatic digests are required; the analyte is a single denatured unfolded protein molecule and a monomer is a residue.

Normally a blockade current resolution able to discriminate among the standard 20 residue types would be required. Such resolution is unattainable in practice, especially with noise present. This is mitigated somewhat if a four-way division of the 20 amino acids based on volume is used [13].

The method presented here makes the resolution problem manageable by reducing the measured number of blockade current signal levels from 20 to two. Subsequences in the resulting binary sequence that can uniquely identify the protein in a binary-coded proteome database are then found. As noted earlier and demonstrated below, computational analysis of a sample proteome (*H. pylori*) using amino acid volume data [20] indicates that almost all of the proteins therein can be identified with a confidence level exceeding 90%.

The proposed method is analyzed computationally before considering implementation issues.

## 3. Computational analysis and results

There are three steps: 1) Divide the set of amino acids into two ordered subsets S_1_ and S_2_; 2) Recode the primary sequences in a proteome with a binary code based on this two-way partition; 3) For every protein in the proteome find one or more subsequences in its binary-coded primary sequence that identify the protein uniquely.

Table 1 shows the standard 20 amino acids grouped by volume into two subsets and shaded by group. The dividing line is between P and V so that the subsets have roughly similar sizes (8 and 12) and the difference between the volumes of P and V is maximal. The amino acids are coded as follows:

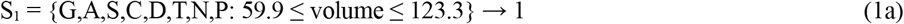

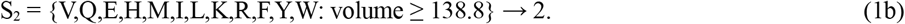

**Table 1.** Amino acids sorted by volume. AA = amino acid. Mean and S.D. (Standard Deviation) in 10^−3^ nm^3^. Data from [20].

| AA | Mean | S.D. | AA | Mean | S.D. | AA | Mean | S.D. | AA | Mean | S.D. |
| --- | --- | --- | --- | --- | --- | --- | --- | --- | --- | --- | --- |
| G | 59.9 | 2.2 | T | 118.3 | 2.3 | Q | 145.1 | 5.1 | K | 172.7 | 5.9 |
| A | 87.8 | 2.3 | N | 120.1 | 4.1 | H | 156.3 | 6.1 | R | 188.2 | 9.6 |
| S | 91.7 | 1.8 | P | 123.3 | 1.8 | M | 165.2 | 1.8 | F | 189.7 | 7.4 |
| C | 105.4 | 5 | V | 138.8 | 3.6 | I | 166.1 | 3.4 | Y | 191.2 | 8 |
| D | 115.4 | 2.2 | E | 140.9 | 5.3 | L | 168 | 4.3 | W | 227.9 | 3.8 |

It was recently shown experimentally that a sub-nanometer-diameter nanopore can measure residue volume with a resolution of 0.07 nm^3^ [23]. Such a level of resolution is sufficient to determine that a residue in a translocating protein belongs to S_1_ or to S_2_. In what follows, residue volume is used as a proxy for the blockade current.

From Table 1, a blockade current threshold T_2_ corresponding to a volume of ~0.13 nm^3^ can distinguish residues in S_1_ from those in S_2_. To distinguish blockades due to residues in S_1_ from the open pore current a second threshold T_1_ corresponding to a volume of ~0.05 nm^3^ is set. Thus if the measured volume of a residue is between T_1_ and T_2_ the output is ‘1’; if > T_2_ the output is ‘2’.

Errors in the resulting binary codes for subsequences of different lengths can be estimated for different volume thresholds. Assuming residue volumes to be normally distributed with mean μ and standard deviation σ, the errors can be computed with the normal (Gaussian) error function. Fig. 1 shows normal distributions of the 20 amino acids based on data from Table 1.

**Fig. 1.**
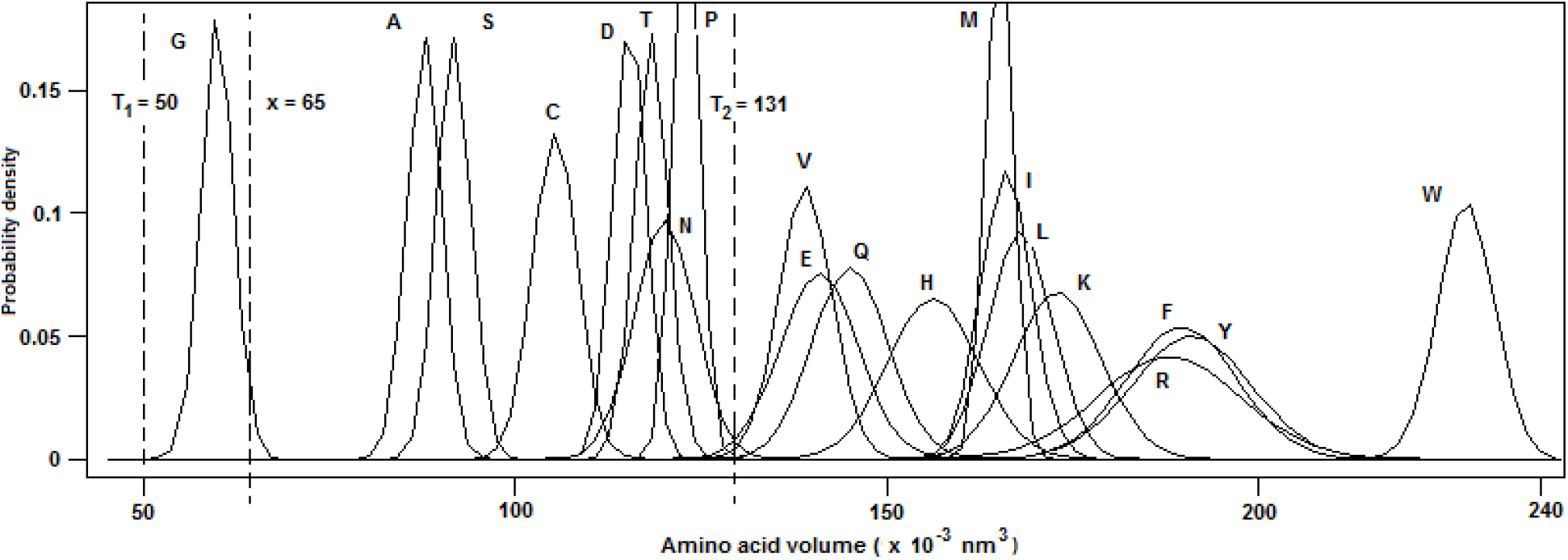
Normal distributions of amino acid volumes

Let the mean volume and standard deviation for amino acid aa be μ_aa_ and σ_aa_ (Table 1). Let the error in reading the volume of amino acid aa be e_aa_(T_1_, T_2_). If F(x; μ, σ) is the cumulative normal distribution function with mean μ and standard deviation σ, the errors for the 20 amino acids are given by

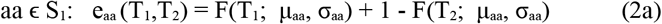

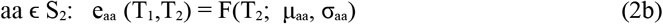

Assuming that blockades due to successive residues are independent the probability that the measured binary code for a protein sequence **X** = X_1_ X_2_…X_n_, where X_i_ is one of the 20 amino acids, is correct (that is, its confidence level) is given by

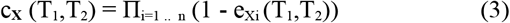

The proteome of the gut bacterium *H. pylori* (Uniprot id UP000000210, 1553 sequences, www.uniprot.org) is used as an example. Supplementary File 1 contains the complete set of protein sequences recoded in binary according to the binary code in Equation 1. Fig. 2 (symbol ♦) shows the percentage of binary-coded subsequences with confidence level > 90% versus subsequence length.

**Fig. 2.**
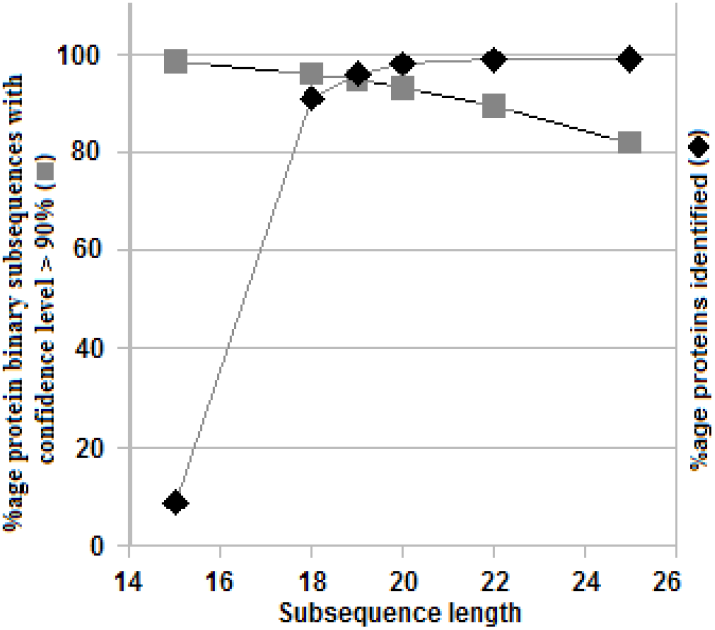
Percentage of protein subsequences in H. pylori whose volume-based binary codes have a confidence level > 90% vs subsequence length (■). Percentage of proteins identified uniquely from subsequences (♦). Thresholds: T_1_ = 0.05 nm^3^, T_2_ = 0.13 nm^3^.

Subsequences from every protein in a proteome are exhaustively compared with every other protein to determine if they uniquely identify their container proteins. To reduce computation time candidate subsequences used are spaced Δ = 5 residues apart. The percentages of proteins identified are given for subsequence length L = 15, 18, 19, 20, 22, and 25 in Fig. 2 (symbol ■). The number of proteins identified goes from 8.69% with L = 15 to ~98% with L = 20 and 99.1% with L = 25. L > 20 yields diminishing returns; gains from reducing Δ are minimal. L = 20 is an optimal length for subsequences.

A complete list of protein identifying subsequences for all the proteins of *H. pylori* is given in Supplementary File 2.

### 3.1 Reduced false detection rate

A significant majority of proteins in the *H. pylori* proteome have a large number of identifying subsequences; this reduces the false detection rate (FDR) considerably. Fig. 3 shows the distribution of the number of proteins vs the number of subsequences of length 20 that identify them. Of the 1553 proteins in the proteome 1501 proteins have more than one identifier, 23 one, 29 none, and 304 more than 40.

**Fig. 3.**
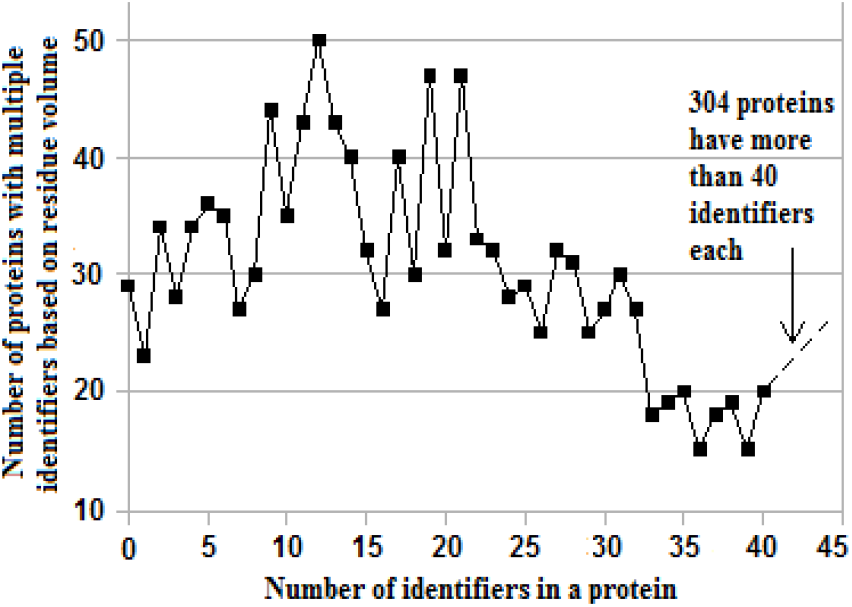
Number of proteins vs number of protein identifying subsequences of length 20 in a protein in H. pylori (1553 sequences)

### 3.2 Reducing detector bandwidth

Normally an analyte like a protein molecule translocates through the pore rapidly (diffusion is the major cause, although the electrophoretic force due to the applied potential and other factors also play a role [24]). This makes it difficult for a detector with insufficient bandwidth to detect changes in the blockade current level. (The limit at present is ~1 MHz [25].) One way to reduce the required bandwidth is to compute an average over the raw pore current with hardware or software. This is essentially a smoothing technique that also leads to a decrease in the number of proteins that can be identified. It is shown next that with this approach the required bandwidth can be reduced by a factor of 10 or more with only a 10% decrease in the number of proteins identified.

Let the detector time resolution be τ, the corresponding detector bandwidth B = 1/2τ. With this up to L blockade pulses can be identified in a pore current signal interval of width 2Lτ. With a bandwidth of B/L (or equivalently a time resolution of Lτ), a pore current pulse of width Lτ can be detected and an average over this interval computed. The result is a sequence of average signal values from which in principle the binary-coded primary sequence can be extracted. Alternatively the average over an interval of width Lτ can be approximated by the number of 2’s (or 1’s) in that interval. Let N_2_(i) be the number of 2’s in the interval [(2iLτ, (2i+1)Lτ], i = 0,1,2,… The sequence of (continuous) average values over alternating intervals of width Lτ can now be approximated by the sequence of integers {N_2_(i); i = 0,1,2,…} corresponding to the number of 2’s in [0, Lτ], [2Lτ, 3Lτ], [4Lτ, 5Lτ], etc. Since 0 ≤ N_2_(i) ≤ L, this is a sequence of numbers in base L+1. Subsequences of length K from the sequence, that is, N_2_(j) N_2_(j+1)…N_2_(j+K-1), j ≥ 0, can now be used as identifiers if they identify the parent protein uniquely. ESM File 2 has two examples showing how this works.

A second sequence {N_2_’(i); i = 0,1,2,…} can be defined with sample intervals [Lτ, 2Lτ], [3Lτ, 4Lτ], [5Lτ, 6Lτ],…(For example, in protein 0 above this gives _4_3_1_5_3_5, where _ stands for the sample interval [2k, (2k+1)Lτ], k ≥ 0). By counting 1’s instead of 2’s two more sequences {N_1_(i); i = 0,1,2,…} and {N_1_’(i); i = 0,1,2,…} can be defined. From these four sequences subsequences of length K can be examined to determine if they are identifiers. The total number of identified proteins is then the cardinality of the union of the four sets of proteins identified by subsequences of the four sequences {N_2_(i); i = 0,1,2,…}, {N_2_’(i); i = 0,1,2,…}, {N_1_(i); i = 0,1,2,…}, and {N_1_’(i); i = 0,1,2,…}. More generally sample intervals over which averaging is done can start anywhere along the primary sequence.

Table 2 gives the results for different values of L and K; it shows a trade off between the bandwidth and the number of proteins identified. With L = 5 and K = 8 the bandwidth is reduced by a factor of 10 while the number of proteins identified in *H. pylori* falls from 1524 (98.13%) to 1372 (88.35%). (This process can be repeated with 1’s instead of 2’s, but the increase is marginal: with L = 5 and K = 8 the number goes up from 1372 to 1375.) ESM File 3 contains a complete list of protein identifiers for *H. pylori* based on averaged subsequences for the case L = 5 and K = 8.

**Table 2.** Bandwidth reduction with averaging. Average over alternating windows of width L (= length of subsequence) is given by number of 2’s in subsequence binary code. Resulting sequence of averages is an (L+1)-ary sequence; an id is an (L+1)-ary subsequence thereof of length K. Data for *H. pylori* (1553 sequences). Id’d = Number of proteins identified in proteome.

|  |  |  |  |  |  |  |  |  |  |  |
| --- | --- | --- | --- | --- | --- | --- | --- | --- | --- | --- |
| L | 5 | 5 | 5 | 6 | 6 | 6 | 6 | 8 | 8 | 8 |
| K | 6 | 8 | 10 | 6 | 8 | 10 | 12 | 6 | 8 | 10 |
| Id'd | 849 | 1372 | 1346 | 1054 | 1337 | 1271 | 1185 | 1197 | 1227 | 1118 |

For other hardware-based approaches to bandwidth reduction see Item 6 in Section 4.

### 3.3 The homopolymer problem

The homopolymer problem refers here to the difficulty in identifying successive residues from the same subset as they generate the same (binary) output value. With a thick (8-10 nm) biological or synthetic pore, multiple (typically 4 to 8) residues are resident in the pore at any time during translocation so that the boundary between two successive values may be hard to detect, although correlations in the measured signal can often provide useful information. Thus in [13] the blockade current was found to correlate well with four contiguous residues.

The averaging technique of Section 3.2 mitigates the homopolymer problem. As averaging is done over a whole interval, the boundary between two successive residues is no longer relevant.

For other solutions based on hardware or software, see Item 7 in Section 4 below.

### 3.4 Quantifying proteins in a mixture

The procedure described here can be used to quantify proteins in a mixture of proteins {M_i_: i = 1, 2,…}, where M_i_ is the number of molecules of the i-th protein. Let 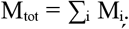. If molecules enter the pore in some random order, then after a sufficiently long run M_i_/M_tot_ can be estimated as 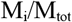 where 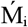 and 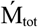 are the measured number of molecules of protein i and the total measured molecules over the run. Quantification time can be reduced significantly by using an array of pores.

## 4. Discussion

Some potential implementation and other relevant issues are now considered.

1. The method described here works on single unbroken protein molecules, no proteolysis is done; it is thus free from the vagaries of the latter [3]. As there is no degradation the sample can be reused/resequenced.
2. Searching for a measured subsequence in the proteome will require both forward and reverse matches because the protein may enter the pore C- or N-terminal first.
3. Equation 3 assumes that successive residues in a protein are independent. This is not true in practice as there are inherent correlations. The latter can be extracted from the pore current signal and used in error correction, this leads to increased reliability. Software used in nanopore-based DNA sequencing routinely uses this kind of information to improve sequencing accuracy [26,27].
4. Depending on the primary sequence a protein may carry only a weak charge so that entry into and/or translocation through the pore may be a problem. One solution [16] to this is to attach a negatively charged carrier molecule like DNA to the protein molecule; another may be based on dielectrophoretic trapping [28].
5. Charged residues on the pore wall tend to interfere with the passage of an analyte when the latter has charged residues. (Seven amino acids, namely D, E, K, R, H, C, and Y, carry a negative or positive charge whose value depends on the pH of the electrolyte.) To resolve this the wall charge can be neutralized in some way. With DNA as the analyte a lipid coat has been shown to do this [29].
6. Hardware solutions to the bandwidth reduction problem of Section 3.2 include use of: a) a room-temperature ionic liquid (RTIL), which is a high viscosity electrolyte that can slow down an analyte by a factor of ~200 [30]; b) an opposing hydraulic pressure field [31]; c) an enzyme (‘unfoldase’) to unfold the protein molecule before it enters into and thus slow its passage through [32] the pore; or d) ligands attached to the protein or the pore [33].
7. The homopolymer problem of Section 3.3 has also been addressed with hardware. If a single atom layer of graphene [34] or molybdenum sulphide (MoS_2_) [30] is used for the membrane, or a biological pore with a narrow constriction (MspA, CsgG) [35,36], or an adapter such as β-cyclodextrin in αHL [37], roughly one residue will be resident in the pore or its constriction or in the adapter during translocation. Software based on a Hidden Markov Model [26] or Viterbi algorithm [27] may also be used to computationally separate successive residues with near identical (binary-coded) blockade levels.
8. This approach also works if residue volume is replaced with residue mass. Following [1] the 20 amino acids are listed here in array form ordered on mass:

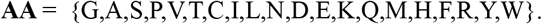

Table 3 shows their division into two subsets of sizes 11 and 9 (shown shaded by subset in the table).

**Table 3.** Amino acids sorted by mass. AA = amino acid. Mass in dalton. Data based on [1].

| AA | G | A | S | P | V | T | C | I | L | N |
| --- | --- | --- | --- | --- | --- | --- | --- | --- | --- | --- |
| Mass | 57.02 | 71.04 | 87.03 | 97.05 | 99.07 | 101.05 | 103.01 | 113.08 | 113.08 | 114.04 |
| AA | D | E | K | Q | M | H | F | R | Y | W |
| Mass | 115.03 | 128.06 | 128.09 | 129.04 | 131.04 | 137.06 | 147.07 | 156.1 | 163.06 | 186.08 |

The subsets are coded with the binary alphabet {1,2}:

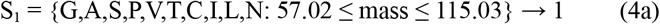

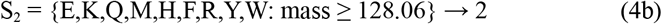

Similar to division by volume, the dividing line is chosen between D and E so that the two sets have roughly similar sizes (11 and 9) and the difference between the masses of D and E is relatively large. The primary sequences in the proteome are then recoded with the mapping given in Equation 4. This leads to a binary-mass-coded proteome sequence database (Supplementary File 3).

The procedure for generating residue-mass-based identifiers for the proteins of *H. pylori* is identical to Step 2 above. Fig. 4 below is the mass counterpart of Fig. 3. In this case 1509 proteins have more than one subsequence identifier of length 20, 15 one, 29 none, and 263 more than 40. As with binary-volume-coding, the identity of most proteins is redundantly encoded in the binary-mass-coded primary sequence. The data behind Fig. 4 are available in Supplementary File 4.

**Fig. 4.**
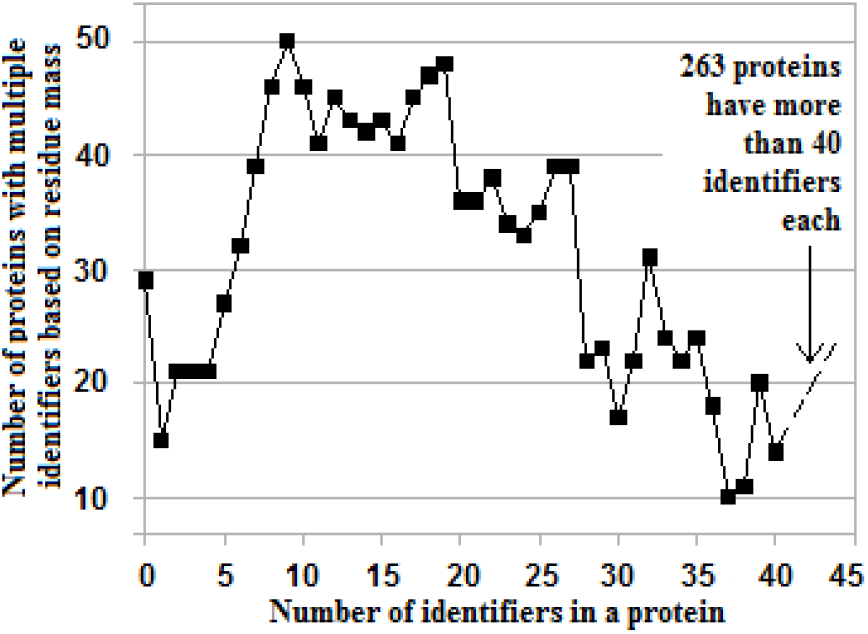
Distribution of number of proteins vs number of protein identifying subsequences of length 20 based on residue mass in a protein in *H. pylori* (1553 sequences).

However, unlike residue volume, which translates to current blockade level in a nanopore, there is no similar measurable behavior for residue mass, which is central to MS. The extraordinary precision with which it is measured by mass spectrometry (MS) [38] in combination with machine language algorithms [39], allows proteins to be sequenced (not just identified) with a high level of confidence. This applies to *de novo* sequencing as well so proteins whose provenance is not known or those designed *de novo* can also be sequenced. An attempt is currently being made to determine if binary-coded residue mass is useful in MS-based studies of proteins, in particular those with post-translational modifications [40].

## 5. Conclusion

Unlike most recent work in protein identification, which is usually aimed at identifying specific single proteins or their variants, the method proposed in this Letter can identify an arbitrary protein in a large set such as a proteome. The availability of sub-nanopores capable of measuring residue volumes with adequate resolution makes the proposed method both feasible and practical. Reducing the number of current blockade levels to be measured from 20 to two effectively removes a major obstacle to the use of nanopores for protein identification. All of this points to the near-term prospect of implementing an easy-to-use hand-held device that can identify and quantify proteins in mixtures while requiring minimal sample preparation without any proteolysis.

**Note**: The file format for Supplementary Files 1 and 2 is given in a preamble to the data. File format information for Supplementary Files 3 and 4 is not included, refer to Files 1 and 2 respectively.

